# Terminal Modification Reprograms Supramolecular Organization of Integrin Ligand Assemblies to Shape Cell Migration

**DOI:** 10.64898/2026.09.29.755244

**Authors:** Ziyi Su, Yanling Lin, Zhu Zhu, Xiaoqing Li, Wenjing Hou, Qizheng Zhang, Yanbin Cai, Jie Gao, Zhimou Yang, Ye Zhang, Xunwu Hu

## Abstract

Precise organization of bioactive ligands at material-cell interfaces is critical for directing integrin-mediated cell behavior, yet tuning supramolecular organization commonly necessitates orchestrated redesign of molecular composition or ligand sequence. Here we establish terminal modification as a minimal, sequence-preserving strategy to reprogram the organization and function of self-assembling integrin ligands. Using FFFIKVAV as a model system, N- and C-terminal modifications generated distinct supramolecular states with markedly different integrin activation and cell migration responses. Notably, stronger intermolecular association did not produce stronger biological activity, as N-terminal acetylation promoted scaffold consolidation while reducing the configurational freedom of the exposed IKVAV ligand and diminishing integrin activation and cell migration. Alongside modulating migration magnitude, terminal chemistry also diversified the cellular programs supporting migration, with the parent peptide and its C-terminally amidated analogue favoring protrusion-dominant and stress-fiber-dominant adhesion–cytoskeletal programs, respectively. These functional states were reproduced when the assemblies were introduced directly to pre-adherent cells, extending their activity beyond preformed material coatings. Our findings identify terminal chemistry as a compact molecular design variable that links supramolecular organization to ligand function while diversifying the biological output of a shared bioactive sequence.

---

Spatial presentation, local clustering, and the nanoscale architecture of integrin ligands^1^ can alter receptor engagement^2^, focal adhesion formation^3, 4^, cytoskeletal organization^5^, and ultimately cell migration^6^. Supramolecular biomaterials provide a particularly useful platform for controlling these parameters because molecular building blocks can self-organize into higher-order structures while presenting bioactive ligands at material-cell interfaces^7–9^. However, manipulating these parameters often requires changes in ligand content^10^, molecular composition^11^, or scaffold design^12^, making it difficult to isolate how supramolecular organization itself contributes to cellular responses^9^. A molecularly minimal strategy that can reconfigure integrin ligand assemblies while preserving the bioactive sequence would therefore provide a powerful means to decouple assembly architecture from ligand identity.

Terminal modification provides a particularly simple means of perturbing molecular interactions within peptide assemblies, with N-terminal acetylation and C-terminal amidation selectively altering terminal charge states and hydrogen-bonding capabilities while preserving the internal amino acid sequence. In short self-assembling peptides, where hierarchical assembly arises from a delicate balance of aromatic, hydrophobic, electrostatic, and hydrogen-bonding interactions^13, 14^, such seemingly minor chemical changes can be amplified into substantial differences in molecular packing and higher-order assembly^15, 16^. This offers an unusual opportunity to reconfigure assembly architecture while holding both the assembling motif and the bioactive ligand sequence constant. Yet whether such minimal chemical changes can propagate from molecular packing to integrin-dependent cellular behavior remains largely unresolved. Establishing this connection would position terminal modification not merely as a chemical capping strategy, but as a molecular design principle for reprogramming bioactive supramolecular assemblies to shape biological responses.

Here, by systematically varying terminal chemistry while preserving both the assembly-driving and integrin-binding sequences, we show that minimal molecular modifications can reprogram supramolecular organization and cell migration. Molecular simulations further link assembly-core consolidation to altered conformational freedom of the exposed ligand, providing a model for how terminal chemistry reshapes the ligand-bearing interface. These distinct assembly states produced divergent patterns of integrin activation and adhesion-cytoskeletal organization, giving rise to different migration-associated cellular programs. Importantly, these phenotypes were recapitulated when the assemblies were introduced directly to pre-adherent cells, extending this design principle beyond preformed substrate coatings and suggesting potential for therapeutic modulation of cell-matrix interactions within pathologically remodeled tissue microenvironments. Together, our findings establish terminal chemistry as a minimal molecular design strategy for directing cell behavior through the supramolecular organization of a shared bioactive ligand.

## RESULTS AND DISCUSSION

### Terminal Modification Reprograms the Supramolecular Organization of Integrin Ligand Assemblies

To determine how minimal terminal chemistry is translated into supramolecular organization, FFFIKVAV^17^ was chosen as a chemically compact yet biologically validated supramolecular ligand platform (Figure 1a). In this architecture, the FFF segment provides a defined assembly-driving element^18^, whereas the laminin-derived IKVAV motif confers integrin-binding activity^19^. Previous work further established that supramolecular organization within this platform can regulate integrin engagement and downstream cellular responses^10, 17^, making FFFIKVAV well suited for tracing how subtle molecular perturbations propagate across scales. N-terminal acetylation and C-terminal amidation were therefore introduced as chemically minimal perturbations that alter terminal charge states and hydrogen-bonding environments while preserving the self-assembling building block and integrin-binding sequence. Accordingly, four terminal variants were generated, comprising FV (H-FFFIKVAV-OH), Ac-FV (Ac-FFFIKVAV-OH), FV-NH₂ (H-FFFIKVAV-NH₂), and Ac-FV-NH₂ (Ac-FFFIKVAV-NH₂) (Figure 1b). A scrambled-sequence analogue, sFV (H-FFFVKIAV-OH), which retains self-assembly but lacks integrin-ligand recognition, was included as a control (Figure S1).

**Figure 1.**
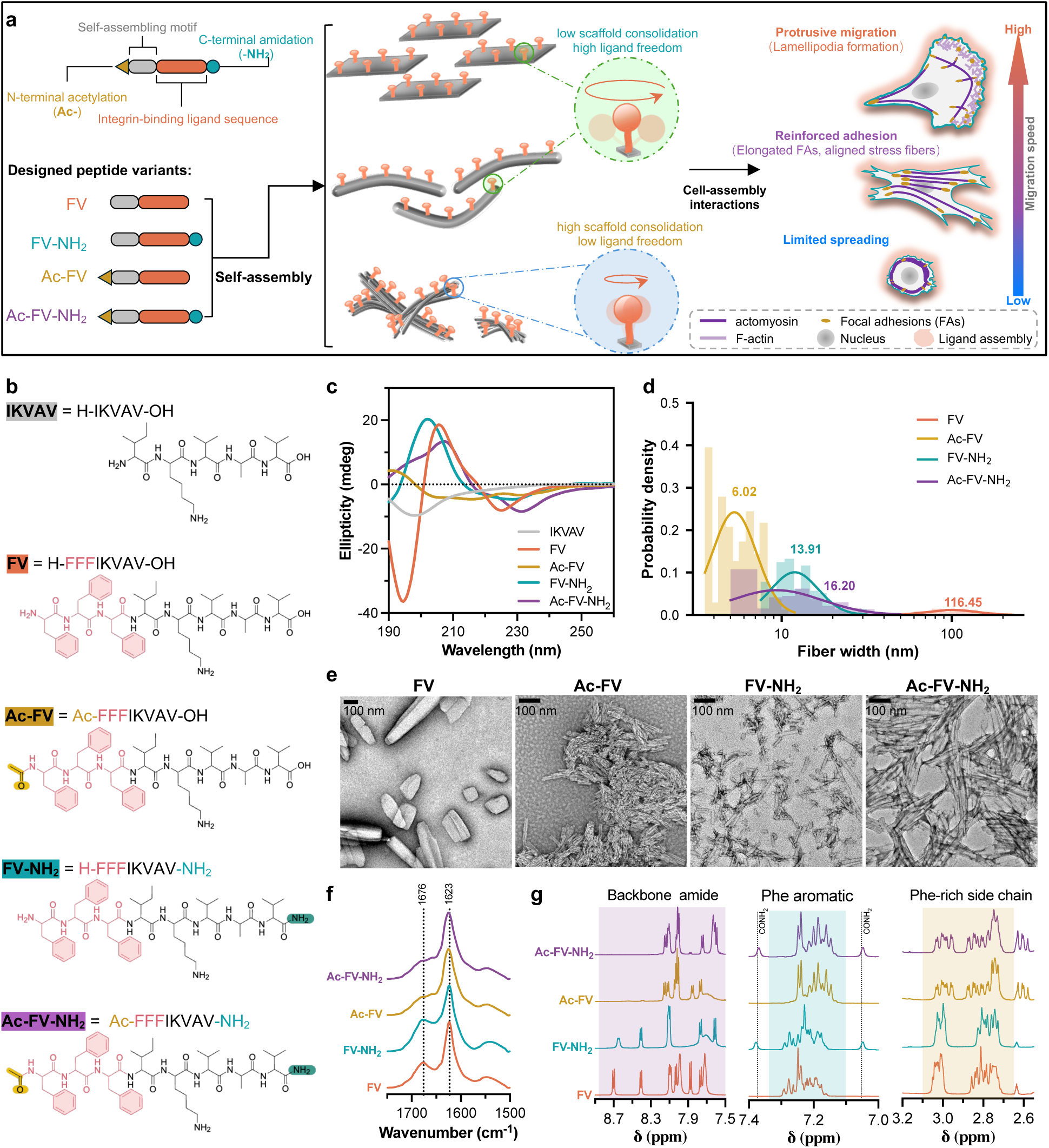
Terminal modification regulates the supramolecular organization of integrin ligand assemblies. **a** Schematic illustration of terminal modification of integrin ligands to regulate supramolecular organization and cell migration. **b** Chemical structures of IKVAV, FV, Ac-FV, FV-NH₂, and Ac-FV-NH₂. FV denotes H-FFFIKVAV-OH, with Ac and NH₂ indicating N-terminal acetylation and C-terminal amidation, respectively. **c** Circular dichroism (CD) spectra of IKVAV and the four peptide variants in water at 200 μM. **d-e** Width distributions (**d**) and representative transmission electron microscopy (TEM) images (**e**) of FV, Ac-FV, FV-NH₂, and Ac-FV-NH₂ assemblies prepared in water at 200 μM. Numbers indicate the mean widths. Scale bars represent 100 nm. **f** Attenuated total reflectance Fourier-transform infrared (ATR-FTIR) spectra of peptide assemblies lyophilized from 200 μM solutions in D₂O. Dashed lines indicate 1676 and 1623 cm⁻¹. **g** Selected regions of the ¹H nuclear magnetic resonance (NMR) spectra of the four peptides, showing the backbone amide, Phe aromatic, and Phe-rich side-chain regions. Dashed lines indicate terminal carboxamide (–CONH₂) resonances.

Circular dichroism (CD) spectroscopy revealed pronounced terminal-dependent changes in supramolecular ordering (Figure 1c and Figure S2). FV exhibited a positive band near 205.5 nm and a negative band around 225 nm, consistent with a red-shifted, noncanonical β-sheet-associated CD signature with contributions from aromatic interactions among the FFF residues^20, 21^. Ac-FV showed a markedly attenuated and blue-shifted spectrum, whereas the amidated variants retained pronounced long-wavelength negative bands near 229-232 nm. These spectral changes are consistent with terminal-dependent reorganization of β-sheet-associated supramolecular ordering. This divergence was further amplified at the morphological level. TEM showed that, unlike IKVAV alone, all FFF-containing variants formed fibrillar assemblies, but with markedly different lateral organization (Figure 1d-e and Figure S3). FV formed broad, laterally associated structures (116.45 nm), whereas either N-terminal acetylation or C-terminal amidation produced much thinner fibrils, measuring 6.02 nm for Ac-FV and 13.91 nm for FV-NH₂. Notably, amidation within the N-acetylated background increased the width to 16.20 nm for Ac-FV-NH₂, indicating a context-dependent interplay between N- and C-terminal chemistry in directing higher-order assembly.

To further examine the molecular consequences of terminal modification, FTIR and ^1^H NMR spectroscopy were used to probe backbone organization and local chemical environments (Figure 1f-g and Figure S4). FTIR showed that all variants retained a dominant amide-I band at 1622-1626 cm⁻¹, indicating preservation of a β-sheet-associated hydrogen-bonded framework^14, 20, 22^ despite their distinct fibrillar morphologies. A higher-wavenumber component at 1670-1676 cm⁻¹ was evident in the non-acetylated variants but substantially attenuated upon N-terminal acetylation, consistent with reorganization of β-sheet-associated packing and hydrogen-bonding environments^14, 23^. ^1^H NMR revealed a corresponding terminal-dependent grouping across the backbone amide, Phe aromatic, and side-chain regions dominated by Phe and Lys contributions: FV and FV-NH₂ displayed closely related spectral fingerprints, whereas Ac-FV and Ac-FV-NH₂ showed greater intragroup similarity. Thus, these spectroscopic signatures distinguished acetylated from non-acetylated variants at the molecular level, while the divergent fibrillar morphologies revealed context-dependent contributions from C-terminal chemistry to higher-order organization.

### Terminal Chemistry Modulates the Physicochemical Properties of Integrin Ligand Assemblies

To examine the physicochemical environments associated with these distinct supramolecular structures, we measured concentration-dependent Nile Red fluorescence^24^. The estimated critical aggregation concentrations (CACs) were 184.4 μM for FV, 90.0 μM for FV-NH₂, 11.1 μM for Ac-FV, and 32.7 μM for Ac-FV-NH₂ (Figure 2a). Both acetylated variants generated Nile Red-accessible hydrophobic environments at substantially lower concentrations than their non-acetylated counterparts. C-terminal amidation shifted the estimated CAC in opposite directions, lowering it for FV but increasing it for Ac-FV, indicating that its effects on aggregation depend on N-terminal chemistry. We next monitored Nile Red fluorescence during heating to 55 °C and subsequent return to room temperature to assess the thermal response of these hydrophobic environments^25, 26^. Fluorescence decreased during heating in all four systems. After cooling, FV and FV-NH₂ recovered to approximately 90-100% of their initial signals, whereas Ac-FV and Ac-FV-NH₂ recovered to approximately 70% (Figure 2b). Thus, terminal chemistry influenced both the onset of detectable hydrophobic association and the recovery of the Nile Red-reported microenvironment following thermal perturbation.

**Figure 2.**
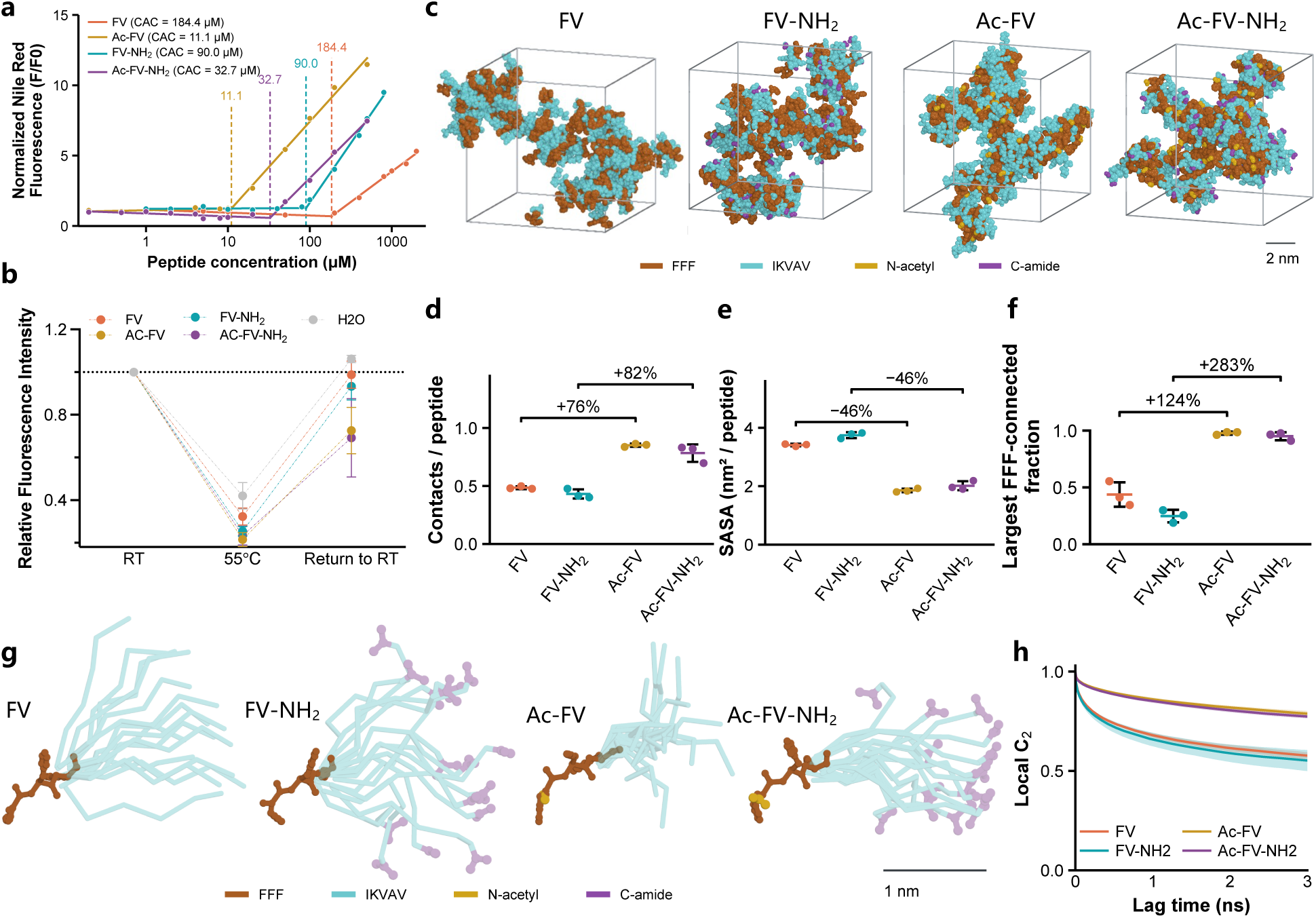
Terminal Chemistry Modulates the Physicochemical Properties of Integrin Ligand Assemblies. **a** Nile Red fluorescence of FV, FV-NH₂, Ac-FV, and Ac-FV-NH₂ at different peptide concentrations. Dashed lines indicate the calculated critical aggregation concentrations (CACs), determined from the intersections of linear fits to the low- and high-concentration regimes. **b** Relative Nile Red fluorescence of peptide assemblies at room temperature (RT), upon heating to 55 °C, and after return to RT. Fluorescence intensities were normalized to the corresponding initial values at RT. Data are presented as mean ± s.d. **c** Snapshots of peptide assemblies from all-atom molecular dynamics (MD) simulations of FV, FV-NH₂, Ac-FV, and Ac-FV-NH₂. FFF, IKVAV, N-acetyl, and C-amide moieties are shown in brown, cyan, yellow, and purple, respectively. Scale bar represents 2 nm. **d-f** Quantification of interpeptide aromatic contacts per peptide (**d**), solvent-accessible surface area (SASA) per peptide (**e**), and the fraction of peptides contained in the largest FFF-connected cluster (**f**) from MD simulations. Each point represents one independent simulation replica (n = 3). Numbers above brackets indicate the relative percentage changes between the indicated groups. **g** Superposition of peptide conformations sampled from MD trajectories after alignment of the FFF motif, showing the spatial distributions of the IKVAV segment and terminal modifications. FFF, IKVAV, N-acetyl, and C-amide moieties are colored as in **c**. Scale bar represents 1 nm. **h** Local second-order orientational autocorrelation functions (C₂) calculated from the MD trajectories for FV, FV-NH₂, Ac-FV, and Ac-FV-NH₂. At least three independent experiments were performed for the experimental measurements.

We used molecular dynamics (MD) simulations to examine the FFF-core organization associated with these differences in aggregation behavior (Figure 2c and Figure S5). Analysis of the 25–50 ns window showed that Ac-FV and Ac-FV-NH₂ exhibited 76% and 82% more aromatic contacts per peptide than FV and FV-NH₂, respectively (Figure 2d). These increases were accompanied by an approximately 46% reduction in the solvent-accessible surface area (SASA) of the FFF core in both comparisons (Figure 2e). The fraction of peptides incorporated into the largest FFF-connected network increased by 124% and 283%, respectively, indicating more extensive connectivity among the assembly-driving segments (Figure 2f). The acetylation-dependent grouping was maintained across independent trajectories, full- and late-window analyses, and alternative connectivity definitions (Figures S6-S8). The increased aromatic association, greater interpeptide connectivity, and reduced core solvent exposure support a more consolidated FFF-rich core in the simulated acetylated assemblies. This molecular organization is consistent with their lower experimentally estimated CACs.

The differences in FFF-core organization prompted us to examine the configurational behavior and solvent exposure of the attached IKVAV motifs. Superimposed configurations following FFF alignment illustrated the conformations sampled by the IKVAV segments (Figure 2g). Scaffold-relative root-mean-square fluctuations (RMSF) of the IKVAV decreased by approximately 27% upon acetylation of FV and by 26% upon acetylation of FV-NH₂ (Figure S9). The local orientational correlation function, C₂, also retained higher values in both acetylated assemblies, indicating more persistent IKVAV orientations relative to the scaffold (Figure 2h). This distinction was reproduced in full- and late-window analyses (Figure S10). Total IKVAV SASA decreased by 0.6% and 9.2% in the corresponding comparisons, showing that the changes in ligand motion were accompanied by comparatively modest changes in overall solvent exposure (Figure S9). Together, these findings support a model in which consolidation of the assembly-driving core is associated with reduced configurational freedom of solvent-exposed IKVAV, connecting terminal chemistry to the organization of both the scaffold and the ligand-bearing interface. Such differences in conformational sampling may influence the ability of exposed IKVAV motifs to adopt receptor-compatible conformations, motivating examination of β1-integrin activation and subsequent cellular responses^2, 3^.

### Supramolecular Organization Shapes Integrin Activation and Cell Migration

Extravillous trophoblast migration and invasion during early placentation^27^ are actively regulated by integrin–ligand interactions with the maternal extracellular matrix^28^. We therefore selected HTR-8/SVneo cells, a first-trimester human trophoblast-derived cell line^29^, to examine how the supramolecular organization of integrin ligands influences cell adhesion and migration on peptide-coated substrates. Based on our previous experience, a coating density of 66.7 nmol cm⁻² was generated to achieve a nearly continuous peptide layer across the substrate^10^. Coating coverage and morphology were assessed to verify the interfaces presented to cells. Congo Red staining showed approximately 90% coverage by the four variants, with little detectable deposition of IKVAV, while atomic force microscopy revealed fibrillar features consistent with TEM observations (Figure S11). We first quantified early cell attachment to determine how effectively cells established initial contacts with the peptide-coated substrates before subsequent spreading and migration. All four terminal variants increased cell attachment relative to the control, with FV and FV-NH₂ producing the highest cell densities, whereas IKVAV and the scrambled analogue sFV did not show significant enhancement (Figure S12). Given the integrin-binding activity of IKVAV, we used FV to validate integrin involvement in attachment. Treatment with the blocking antibody P5D2 reduced FV-mediated attachment from 355.0 to 150.7 cells mm⁻², while control IgG had little effect (Figure S13). These results confirm integrin involvement in FV-mediated attachment and establish terminal-dependent differences in initial cell-material engagement.

To capture the initial development of adhesions following attachment, we selected 1 h after seeding as an early time point for assessing spreading, integrin activation, and adhesion-site organization^30^. At this early stage, FV and FV-NH₂ produced the largest increases in cell spreading, whereas the acetylated variants remained substantially less spread (Figure 3a, b and Figure S14). To relate these spreading differences to receptor engagement, we assessed β1-integrin activation. FV and FV-NH₂ elicited the strongest activation of β1 integrin, whereas Ac-FV showed only a weak response; C-terminal amidation partially restored integrin activation in Ac-FV-NH₂, although the response remained markedly below that of the non-acetylated assemblies (Figure 3c, d and Figure S15). Because activated integrins are subsequently organized into adhesion complexes that couple the extracellular substrate to the actin cytoskeleton^5, 31^, we next examined the spatial association of activated integrin with paxillin to assess adhesion-site organization. FV and FV-NH₂ showed pronounced accumulation and spatial colocalization of activated integrin and paxillin at adhesion sites (Figure 3e and Figure S16). In contrast, Ac-FV showed weak and poorly organized integrin–paxillin association, whereas C-terminal amidation partially restored this organization in Ac-FV-NH₂.

**Figure 3.**
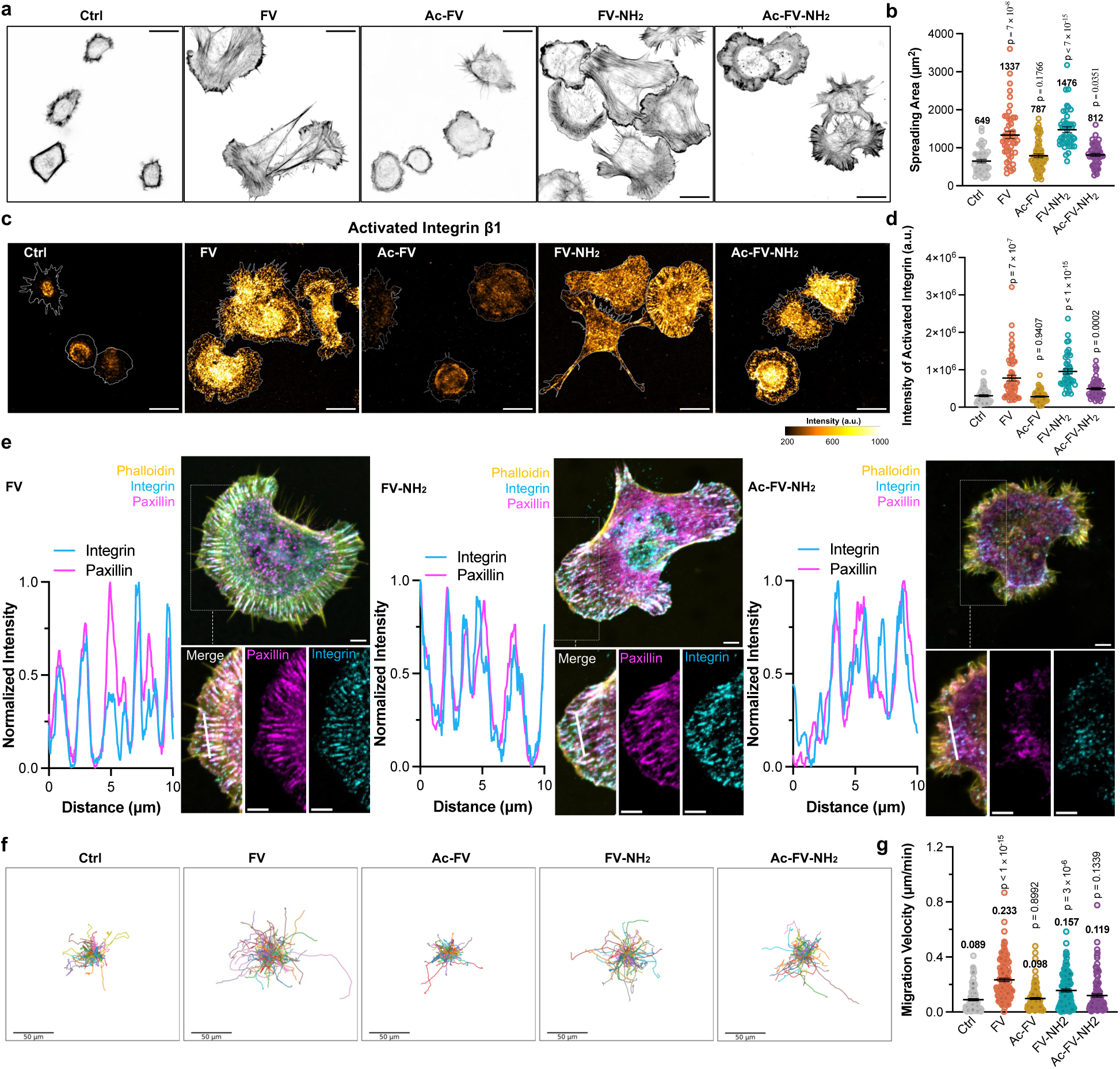
Supramolecular organization of integrin ligand assemblies regulates cell spreading, integrin activation, and migration. **a, b** Representative phalloidin staining images showing F-actin organization (**a**) and quantification of cell spreading area (**b**) of HTR-8/SVneo cells on Ctrl, FV-, Ac-FV-, FV-NH₂-, and Ac-FV-NH₂-coated substrates after 1 h of cell attachment. Scale bar represents 20 μm. **c, d** Representative immunofluorescence images of activated β1 integrin (**c**) and quantification of activated β1 integrin fluorescence intensity (**d**) after 1 h of cell attachment under the indicated conditions. Scale bar represents 20 μm. **e** Representative fluorescence images of F-actin, activated β1 integrin, and paxillin in cells on FV-, FV-NH₂-, and Ac-FV-NH₂-coated substrates after 1 h of cell attachment. Enlarged regions show the merged images and the corresponding paxillin and integrin channels. Normalized fluorescence intensity profiles of integrin and paxillin were measured along the indicated white lines. Scale bar represents 5 μm. **f, g** Migration trajectories (**f**) and migration velocities (**g**) of HTR-8/SVneo cells. Cells were allowed to adhere to the indicated substrates for 12 h before imaging and were then tracked every 6 min for 2 h. Scale bar represents 50 μm. Error bars represent SEM. Brown–Forsythe and Welch ANOVA followed by multiple-comparisons testing was used for statistical analysis. P values are shown in the plots. At least three independent experiments were performed.

We next asked whether the distinct adhesive states translated into effective cell migration. Random single-cell migration was assessed after 12 h of attachment by tracking cells for 2 h at 6-min intervals. FV and FV-NH₂ increased migration velocity to 0.233 and 0.157 μm min⁻¹, respectively, compared with 0.089 μm min⁻¹ in the control, whereas neither acetylated variant showed a significant increase (Figure 3f-g). Previous work showed that Ca²⁺-mediated crosslinking reduced molecular motion in ligand-presenting supramolecular fibers and altered integrin signaling^32^. Motivated by this approach, we used Ca²⁺ as a sequence-preserving perturbation of FV and Ac-FV, which retain free C-terminal carboxyl groups but differ in N-terminal acetylation. Ca²⁺ was introduced during peptide dissolution and assembly, while coated substrates were washed with PBS before cell seeding to minimize direct cellular effects. Ca²⁺ reduced migration velocity on FV from 0.247 to 0.172 μm min⁻¹, whereas Ac-FV showed no corresponding reduction (0.073 to 0.130 μm min⁻¹) (Figure S17).

### Distinct Integrin Ligand Assemblies Engage Divergent Adhesion-Cytoskeletal Programs

Because protrusion, adhesion, traction, and rear retraction can be supported by distinct spatial organizations of the actin cytoskeleton and focal adhesions (FAs)^33, 34^, we examined whether the enhanced migration on FV and FV-NH₂ reflected different cytoskeletal states. After 12 h of attachment, FV induced broad, actin-rich lamellipodia with extensive peripheral ruffling, whereas FV-NH₂ produced elongated cells dominated by thick, aligned stress fibers spanning the cell body (Figure 4a). Both variants increased spreading relative to Ctrl, while IKVAV and sFV did not (Figure 4b and Figure S18). Paxillin staining further separated the two phenotypes, with FV displaying numerous small adhesions across the protrusive lamellipodia and larger adhesions extending inward across the ventral surface, whereas FV-NH₂ showed prominent streak-like FAs aligned with longitudinal actin bundles (Figure 4c and Figure S19). These observations define two distinct migration-associated adhesion-cytoskeletal states, with FV favoring a lamellipodia-rich protrusive program and FV-NH₂ favoring a stress-fiber-dominated contractile program organized around aligned actin bundles and elongated FAs.

**Figure 4.**
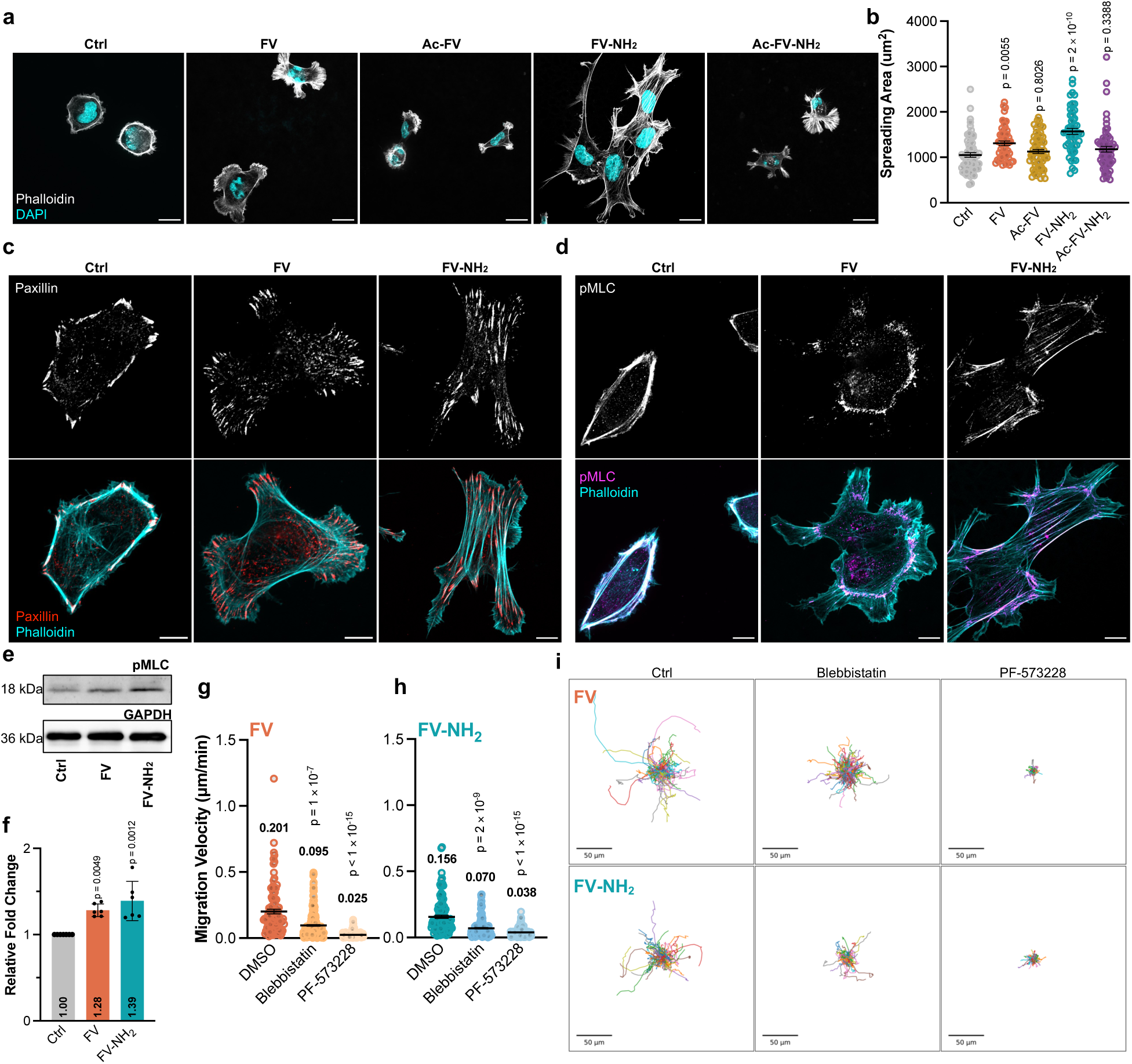
FV and FV-NH₂ induce distinct adhesion–cytoskeletal states associated with enhanced cell migration. **a, b** Representative phalloidin staining images showing F-actin organization (**a**) and quantification of cell spreading area (**b**) of HTR-8/SVneo cells on Ctrl, FV-, Ac-FV-, FV-NH₂-, and Ac-FV-NH₂-coated substrates after 12 h of cell attachment. Nuclei were stained with DAPI. Scale bars represent 20 μm. **c** Representative immunofluorescence images of paxillin and F-actin in HTR-8/SVneo cells on Ctrl, FV-, and FV-NH₂-coated substrates after 12 h of cell attachment. **d** Representative immunofluorescence images of phosphorylated myosin light chain 2 (pMLC) and F-actin under the same conditions. Scale bars in **c, d** represent 10 μm. **e-f** Western blot analysis of pMLC and GAPDH (**e**) and quantification of pMLC expression normalized to GAPDH (**f**) in cells on Ctrl, FV-, and FV-NH₂-coated substrates after 12 h of cell attachment. **g, h** Migration velocities of cells on FV- (**g**) and FV-NH₂-coated (**h**) substrates treated with DMSO, 20 μM blebbistatin, or 10 μM PF-573228. Cells were allowed to adhere for 11 h and treated with the indicated inhibitors for 1 h before migration tracking. **i** Representative migration trajectories under the same conditions as in **g, h**. Cells were tracked every 6 min for 2 h. Scale bars represent 50 μm. Error bars represent SEM. Brown–Forsythe and Welch ANOVA followed by multiple-comparisons testing was used for statistical analysis. Each treatment group was compared with its corresponding control group. P values are shown in the plots. At least three independent experiments were performed.

These distinct FA-actin architectures were accompanied by different patterns of myosin II activity^10, 34^. Phosphorylated myosin light chain 2 (pMLC), a marker of activated myosin II, showed markedly different spatial distributions on FV and FV-NH₂. On FV, pMLC accumulated predominantly in the lamella immediately behind the broad lamellipodium, while the protrusive edge remained F-actin-rich and relatively pMLC-poor. This spatial pattern is consistent with the separation of an actin-rich protrusive edge^10, 33^ from a more contractile lamellar region that supports traction and cell-body translocation^34, 35^. By contrast, FV-NH₂ displayed pMLC along thick, aligned stress fibers traversing the cell body, characteristic of a stress-fiber-based contractile architecture^34, 35^ in which myosin-generated tension is transmitted along adhesion-coupled actin bundles to support traction and polarized movement^36^ (Figure 4d and Figure S20). Immunoblotting further showed increased overall pMLC abundance in both states, reaching 1.28-fold on FV and 1.39-fold on FV-NH₂ relative to Ctrl (Figure 4e-f). Thus, both states engaged myosin II, but its spatial deployment differed markedly between the protrusion-dominant FV and stress-fiber-dominant FV-NH₂ migration modes.

Having established that FV and FV-NH₂ organize myosin II activity into distinct spatial patterns, we next tested whether adhesion-actomyosin coupling^36^ was functionally required for their enhanced migration. The myosin II ATPase inhibitor blebbistatin^37^ reduced migration velocity from 0.201 to 0.095 μm min⁻¹ on FV and from 0.156 to 0.070 μm min⁻¹ on FV-NH₂, while FAK inhibition with PF-573228^38^ produced a stronger suppression to 0.025 and 0.038 μm min⁻¹, respectively, accompanied by markedly restricted trajectories (Figure 4g-i). Neither inhibitor produced a comparable suppression on the control substrate (Figure S21). Taken together, these data define two distinct migration programs elicited by the terminally modified assemblies. Despite their difference in adhesion-cytoskeletal organization, both migration modes required FAK-dependent adhesion signaling and myosin II activity to support productive migration.

### Supramolecular Integrin Ligands Reprogram Cellular Interfaces Beyond Surface Coating

Previous studies showed that self-assembling integrin ligands introduced directly into cell culture can form membrane-associated fibrillar assemblies^10^, engage apical integrins^17, 39^, and remodel adhesion and cytoskeletal organization to regulate cell migration without a preformed ligand-coated substrate^10^. We therefore examined whether the terminally modified assemblies could reproduce their distinct migration-associated states when introduced directly to pre-adherent cells. In the post-adhesion model, cells were allowed to adhere for 12 h before the medium was replaced with peptide-containing medium. Both FV and FV-NH₂ showed good cytocompatibility across the concentrations relevant to subsequent experiments (Figure S22). Concentration-dependent analysis showed that FV enhanced both random single-cell migration and wound closure, with the strongest response at 50 μM (Figure S23). We therefore selected 50 μM for subsequent experiments. At this concentration, FV and FV-NH₂ increased random single-cell migration velocity from 0.101 μm min⁻¹ in Ctrl to 0.241 and 0.175 μm min⁻¹, respectively (Figure 5a-b), while both assemblies also enhanced Transwell efficiency in a complementary Transwell assay (Figure 5c-d). Interestingly, their relative efficacy differed between the two migration contexts, with FV more strongly promoting planar random migration and FV-NH₂ showing the greater Transwell response, consistent with the greater importance of lamellipodial protrusion on planar surfaces and actomyosin contractility during migration through spatially constrained pores^19, 40, 41^.

**Figure 5.**
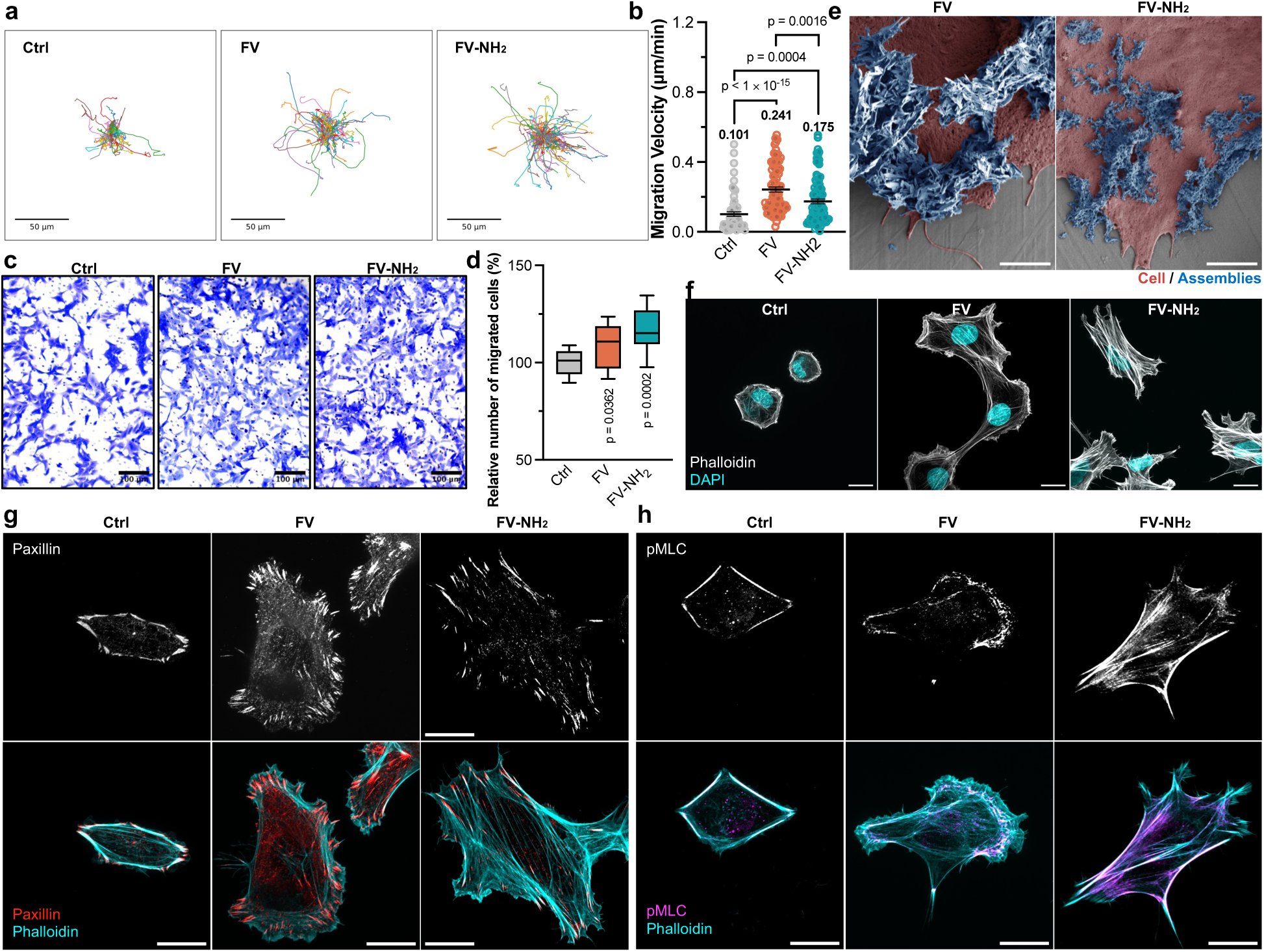
Supramolecular integrin ligands reprogram cellular interfaces beyond surface coating. **a-b** Representative migration trajectories (**a**) and migration velocities (**b**) of HTR-8/SVneo cells treated with 50 μM FV or FV-NH₂. Scale bars represent 50 μm. **c-d** Representative images of Transwell-migrated HTR-8/SVneo cells stained with crystal violet (**c**) and quantification of migrated cells (**d**) in the absence or presence of 50 μM FV or FV-NH₂. Cells were allowed to migrate for 24 h. Scale bars represent 100 μm. **e** Representative false-colored scanning electron microscopy (SEM) images showing interactions between HTR-8/SVneo cells and FV or FV-NH₂ assemblies. Scale bars represent 2 μm. Cells and peptide assemblies are shown in red and blue, respectively. **f** Representative phalloidin staining images showing F-actin organization in the absence or presence of 50 μM FV or FV-NH₂. Scale bars represent 20 μm. Nuclei were stained with DAPI. **g** Representative immunofluorescence images of paxillin and F-actin under the indicated conditions. Scale bars represent 20 μm. **h** Representative immunofluorescence images of phosphorylated myosin light chain 2 (pMLC) and F-actin under the indicated conditions. Scale bars represent 20 μm. Brown–Forsythe and Welch ANOVA followed by multiple-comparisons testing was used for statistical analysis. P values are shown in the plots. At least three independent experiments were performed.

Scanning electron microscopy of cells in this noncoating environment revealed extensive fibrillar assemblies associated with the cell surface for both FV and FV-NH₂ (Figure 5e). FV assemblies preferentially accumulated along cell margins and protrusive regions as broad, densely associated fibrillar structures, whereas FV-NH₂ formed finer and more dispersed fibrillar domains across the cell surface. These distinct nano-morphologies were consistent with the relative structural features observed by TEM and on peptide-coated substrates by AFM, with FV forming broader laterally associated assemblies and FV-NH₂ retaining a finer fibrillar organization. Additional SEM imaging confirmed these characteristic assembly–cell interactions across multiple cells (Figure S24). Consistent with the coating model, FV and FV-NH₂ recapitulated the enhanced spreading phenotype under noncoating conditions, increasing cell spreading area from 671 μm² in Ctrl to 1212 and 1249 μm², respectively, together with corresponding increases in Feret diameter from 44.8 μm to 67.5 and 74.2 μm (Figure S25).

The distinct adhesion-cytoskeletal states observed on coated substrates were also reproduced after direct peptide treatment. FV again induced broadly spread cells with prominent peripheral F-actin and extensive protrusive structures, whereas FV-NH₂ favored a more elongated morphology containing thick, aligned actin bundles (Figure 5f). Paxillin organization showed the same divergence, with FV displaying abundant adhesions across protrusive and ventral regions and FV-NH₂ exhibiting elongated adhesions aligned with longitudinal actin bundles (Figure 5g). A corresponding difference was observed in actomyosin organization, with pMLC enriched behind actin-rich protrusions in FV-treated cells but distributed along prominent stress-fiber-like bundles in FV-NH₂-treated cells (Figure 5h). Together, these results show that adding the assemblies directly to pre-adherent cells recapitulates the protrusion-dominant and stress-fiber-dominant programs observed on coated substrates, demonstrating that terminal-dependent supramolecular organization can reprogram cellular interfaces and migration beyond a preformed surface coating.

## CONCLUSIONS

The design of bioactive supramolecular materials requires coordination between the interactions that build an assembly and the requirements of the ligands presented at its surface. Our results reveal a tension between these objectives within a conserved peptide sequence and identify terminal chemistry as a minimal molecular variable for generating functionally distinct supramolecular interfaces. Here, N-terminal acetylation promoted aggregation at lower concentrations while weakening integrin activation and migration enhancement. Simulations associated acetylation with a more consolidated FFF core and reduced IKVAV conformational freedom, with comparatively modest changes in ligand solvent exposure. These findings support a model in which core packing constrains the configurations available to an exposed ligand. The context-dependent effects of C-terminal amidation further identify terminal-group combinations as an important design variable. Together, these observations suggest that stronger intermolecular association can carry a biological cost. Assembly-core organization and ligand function therefore warrant joint optimization, with terminal chemistry providing a compact route to explore their relationship while preserving both the assembly-driving and bioactive sequences.

Terminal modification also diversified how cells organized migration. FV and FV-NH₂ favored lamellipodia-rich and stress-fiber-dominated adhesion-cytoskeletal states, respectively, while sharing a requirement for focal adhesion kinase and myosin II activity. Their different relative efficacies across migration assays give this distinction functional significance, with FV more strongly promoting planar migration, whereas FV-NH₂ produced the greater Transwell response. Supramolecular ligand design can therefore control not only the magnitude of migration, but also the cellular program through which migration is achieved, allowing material properties to be matched to distinct migration contexts. The reproduction of these states after direct treatment of pre-adherent cells further extends this principle beyond preformed coatings to established cellular interfaces. More broadly, these findings point to a general strategy in which validated bioactive sequences are retained while their supramolecular context is reprogrammed through minimal chemical edits. Extending this principle across other ligand-receptor systems could enable modular nano-bio interfaces that encode distinct cellular responses without requiring redesign of the underlying bioactive motif.

## MATERIALS AND METHODS

### Peptide synthesis

Peptides IKVAV, FFFVKIAV, FFFIKVAV, Ac-FFFIKVAV, and FFFIKVAV-NH₂ were custom-synthesized by solid-phase synthesis with a purity of ≥95% (GL Biochem, China). Ac-FFFIKVAV-NH₂ was custom-synthesized by solid-phase synthesis with a purity of ≥ 95% (GenScript, China). The structures of IKVAV^17^ and FFFIKVAV^10^ were previously reported and validated by NMR and LC-MS in our earlier work. Characterization data for the remaining peptides are provided below.

### Preparation of Peptide Assemblies

Peptides were dissolved in sterile-filtered Milli-Q water at stock concentrations of 10 mM for IKVAV, FV, and FV-NH₂, 5 mM for Ac-FV and Ac-FV-NH₂, and 2 mM for sFV according to their solubility. The solutions were sonicated for 30 min at room temperature, after which the pH was adjusted to 7.0 using 5 M NaOH. Peptide solutions were then incubated overnight at room temperature to allow supramolecular assembly and equilibration. For comparative experiments, the stock solutions were diluted to the same indicated final peptide concentrations using Milli-Q water or cell culture medium, as appropriate.

### Circular Dichroism (CD) Spectroscopy

CD spectra of peptide assemblies were recorded at room temperature using a J-1700 CD spectrometer (JASCO, Japan) under continuous nitrogen purging. Peptide samples were prepared in Milli-Q water at the indicated final concentrations (20–200 μM) and equilibrated overnight at room temperature before being loaded into a 1.0 mm path-length quartz cuvette (Starna, UK). Spectra were collected from 190 to 270 nm in continuous scanning mode. A Milli-Q water blank was measured under identical conditions and subtracted from each spectrum. Each spectrum represents the average of three consecutive scans.

### Fourier Transform Infrared Spectroscopy (FTIR)

Peptide assembly solutions were prepared in deuterated water (D₂O; Innochem, China) at a final concentration of 200 μM and equilibrated overnight at room temperature. The samples were then snap-frozen in liquid nitrogen for 5 min and immediately lyophilized. ATR-FTIR spectra of the lyophilized peptide assemblies were recorded using a Bruker INVENIO-R FTIR spectrometer (Bruker, Germany). Samples were placed directly on a diamond ATR crystal, and spectra were collected at room temperature over 4000–400 cm⁻¹ with a spectral resolution of 4 cm⁻¹. Each spectrum represents the average of 64 scans. Spectral processing was performed using OMNIC software (Thermo Fisher Scientific, USA), including water-vapor subtraction, baseline correction, and min–max normalization over the full spectral range.

### ^1^H Nuclear Magnetic Resonance (NMR) Spectroscopy

Peptide samples (5–10 mg) were transferred to 1.5 mL microcentrifuge tubes, frozen at −80 °C overnight, and subsequently lyophilized to dryness under vacuum. The lyophilized peptides were dissolved in DMSO-d₆ (Adamas; 38114U) at approximately 10 mg mL⁻¹ to obtain clear solutions. Approximately 500 μL of each sample was transferred into a 5 mm NMR tube (Innochem; P9941). ^1^H NMR spectra were recorded at room temperature on a 500 MHz Bruker NMR spectrometer equipped with an Ascend Evo 500 magnet.

### Transmission Electron Microscopy (TEM)

Peptide assembly solutions were prepared at a final concentration of 200 μM and equilibrated overnight at room temperature before imaging. Carbon-coated copper grids were glow-discharged using an SGlow glow discharge system (Beijing Zhongke Chuangyi Technology Co., Ltd., China) to increase surface hydrophilicity. A 10 μL aliquot of each peptide solution was applied to the grid and incubated for 10 s, after which excess liquid was removed with a filter paper. The grids were rinsed three times with 5 μL of distilled water, briefly air-dried, and negatively stained with 5 μL of 1% (w/v) uranyl acetate (SPI Supplies, USA; 02624-AB) for 10 s. Excess stain was removed, and the grids were air-dried before imaging. TEM images were acquired using a JEM-F200 transmission electron microscope (JEOL, Japan) under high-vacuum conditions.

### Nile Red-Based Critical Aggregation Concentration (CAC) Measurement

The critical aggregation concentration (CAC) of the peptides was determined using Nile Red as a fluorescent probe. Nile Red (Sigma-Aldrich, USA) was dissolved in anhydrous methanol to prepare a 2 mM stock solution, stored at −20 °C in the dark, and diluted with Milli-Q water immediately before use. Peptide stock solutions were serially diluted with Milli-Q water to the indicated concentrations. The diluted peptide solutions were sonicated for 10 min and equilibrated overnight at room temperature. Nile Red was then added to each peptide solution at a final concentration of 5 μM. Samples were gently mixed and incubated for 40 min at room temperature in the dark. Fluorescence emission spectra were recorded using a Synergy H1 multimode microplate reader (BioTek, USA), with excitation at 543 nm and emission collected from 580 to 700 nm. The excitation and emission slit widths were both set to 5 nm. Each sample was measured in triplicate. The maximum fluorescence emission intensity was plotted against the logarithm of peptide concentration. The low-concentration and high-concentration regimes were independently fitted by linear regression, and the peptide concentration corresponding to the intersection of the two fitted lines was defined as the CAC. Three independent experiments were performed.

### Thermal Perturbation of Peptide Assemblies Monitored by Nile Red Fluorescence

Peptide assemblies were prepared and incubated with Nile Red using the same procedure described above. Fluorescence was first measured at room temperature to obtain the initial signal. Samples were then heated to 55 °C and maintained at this temperature for 60 min, with fluorescence measurements collected every 10 min. After heating, the samples were returned to room temperature and monitored for an additional 40 min, with measurements collected every 10 min during the recovery period. Fluorescence was measured using a Synergy H1 multimode microplate reader (BioTek, USA) with excitation at 543 nm and emission collected from 580 to 700 nm. Fluorescence intensities were normalized to the corresponding initial values measured before heating.

### Molecular Dynamics (MD) Simulations

Initial peptide structures were generated based on an AlphaFold-predicted FFFIKVAV structure. Terminally modified variants were constructed by modifying the corresponding N- and C-terminal groups in CHARMM-GUI^42^. All simulations were performed at the all-atom level using the CHARMM36-jul2022 force field^43^ in GROMACS 2025.2^44^. For each system, 100 peptide molecules were randomly inserted into a cubic simulation box of 10 × 10 × 10 nm³. The systems were solvated with explicit water and neutralized with Na⁺ and Cl⁻ ions, with an additional NaCl concentration of 0.15 M. Electrostatic interactions were treated using the particle-mesh Ewald (PME) method with a Fourier spacing of 0.12 nm and a real-space cutoff of 1.2 nm; van der Waals interactions were treated with a force-switch modifier between 1.0 and 1.2 nm, without long-range dispersion correction. Bonds involving hydrogen atoms were constrained using the LINCS algorithm. The systems were first energy-minimized using the steepest-descent algorithm until the maximum force fell below 100 kJ mol⁻¹ nm⁻¹. The systems were then sequentially equilibrated for 500 ps under the NVT ensemble (V-rescale thermostat, 303.15 K) and 500 ps under the NPT ensemble (V-rescale thermostat at 303.15 K coupled with a C-rescale barostat at 1 bar, coupling time constant of 5.0 ps, compressibility of 4.5 × 10⁻⁵ bar⁻¹), both using a 1 fs time step and without positional restraints. Three independent production replicas were subsequently simulated for each peptide system under NPT conditions at 303.15 K (Nose-Hoover thermostat) and 1 bar (Parrinello-Rahman barostat) for 50 ns using a 2 fs time step. Periodic boundary conditions were applied throughout all simulation stages. Primary quantitative comparisons were performed using the 25–50 ns portion of each production trajectory. Full-trajectory analyses and five consecutive 10 ns time-block analyses were additionally performed to assess the temporal robustness of the observed differences. Detailed trajectory-analysis procedures are provided in the Supporting Information.

### Preparation of Peptide-Coated Substrates

Peptide stock solutions were diluted with Milli-Q water to a final concentration of 200 μM and equilibrated overnight at room temperature. The equilibrated peptide solutions were then applied to the culture substrates at a nominal peptide loading of 66.7 nmol cm⁻² and dried under vacuum overnight. Before cell seeding or subsequent surface characterization, the coated substrates were washed three times with DPBS to remove loosely associated peptide assemblies.

### Congo Red Staining of Peptide-Coated Substrates

Peptide-coated substrates were stained with Congo Red (Abcam, UK; ab145645) at a 1:1000 dilution for 30 min at room temperature in the dark. The substrates were then washed three times with DPBS, and Z-stack images were acquired using a Nikon AX R confocal microscope system (Nikon, Japan). Z-stacks were projected as maximum-intensity projections (MIPs), and the peptide-coated area was quantified using ImageJ.

### Atomic Force Microscopy (AFM)

For AFM imaging, peptide assemblies were deposited onto freshly cleaved mica at one-quarter of the peptide loading used for cell-coating experiments to allow clear visualization of individual fibrillar structures. The samples were dried under vacuum overnight. AFM images were acquired using a Cypher S atomic force microscope (Oxford Instruments Asylum Research, UK) in tapping mode and analyzed using Nanoscope Analysis software version 1.9 (64 bit).

### Cell Culture

HTR-8/SVneo cells (#SCSP-5203) were obtained from the Cell Bank of the Chinese Academy of Sciences and maintained in Dulbecco’s Modified Eagle Medium (DMEM; Gibco, USA) supplemented with 10% (v/v) fetal bovine serum (FBS; Gibco, USA) and 1% penicillin/streptomycin (Gibco, USA). Cells before passage 8 were used. Cells were routinely tested and confirmed negative for mycoplasma. Unless otherwise specified, all peptide-treatment experiments were performed in DMEM containing 1% FBS to minimize serum-derived interference with cell–material interactions while maintaining cell viability during the experimental period.

### Cell Attachment Assay

Peptide-coated 96-well plates were incubated with 1% bovine serum albumin (BSA; Sigma-Aldrich, USA) for 1 h to block nonspecific cell binding. HTR-8/SVneo cells were resuspended in serum-free DMEM and seeded at 2 × 10⁴ cells per well. For β1-integrin blocking, cells were preincubated with 10 μg mL⁻¹ anti-β1 integrin blocking antibody (clone P5D2; Merck Millipore, USA; MAB1959Z) or 10 μg mL⁻¹ mouse IgG negative-control antibody (Merck Millipore, USA) for 30 min at room temperature before seeding. After 30 min of cell attachment, non-adherent cells were removed, and the wells were gently washed 2 times with DPBS. Adherent cells were stained with Calcein-AM (Beyotime, China) for 20 min and imaged using an EVOS M7000 Imaging System (Thermo Fisher Scientific, USA).

### Cell Seeding on Peptide-Coated Substrates

Following preparation of the peptide-coated substrates, HTR-8/SVneo cells were resuspended and seeded onto the substrates at a density of approximately 7 × 10³ cells cm⁻². Early adhesion-related responses, including cell spreading and integrin-associated adhesion organization, were assessed 1 h after seeding. Unless otherwise specified, downstream experiments, including cell migration, immunostaining, and Western blotting, were performed after 12 h of cell attachment.

### Confocal Microscopy

Confocal imaging was performed using either an AX R Confocal Microscope System (Nikon, Japan) or an X-Light V3 spinning-disk confocal system (CrestOptics, Italy). Cells were seeded on glass-bottom culture dishes (Cellvis, USA; C8SB-1.5H) and subjected to the indicated treatments. After treatment, cells were washed three times with DPBS and fixed with 4% paraformaldehyde in phosphate-buffered solution (Beyotime, China; P0099) for 15 min at room temperature. Cells were then washed three times with DPBS containing 0.3% Triton X-100 (Sigma-Aldrich, USA) for permeabilization. Nuclei and F-actin were stained with NucBlue Fixed Cell Stain ReadyProbes reagent (Invitrogen, USA; 2831256) and ActinGreen 488 (Invitrogen, USA; 2647610), respectively, for 15 min at room temperature. Cells were subsequently washed three times with DPBS containing 0.3% Triton X-100 and imaged by confocal microscopy. Cell morphological features were quantified using ImageJ.

### Immunocytochemistry (ICC)

Cells were subjected to the indicated peptide treatments and subsequently fixed with 4% paraformaldehyde for 30 min at room temperature. After blocking with Immunostaining Blocking Solution (Beyotime, China; P0260) for 1 h, cells were incubated overnight at 4 °C with the indicated primary antibodies diluted in Primary Antibody Dilution Buffer (Beyotime, China; P0277), including anti-paxillin (clone Y113; 1:200; Abcam, UK; ab32084), anti-phospho-myosin light chain 2 (Thr18/Ser19) (1:200; Cell Signaling Technology, USA; 3674S), and anti-activated β1 integrin (clone 9EG7; 1:200; BD Biosciences, USA; 553715). After three washes with DPBS, cells were incubated with the appropriate fluorescent secondary antibodies for 1 h at room temperature in the dark: goat anti-mouse IgG H&L (Alexa Fluor 568; 1:1000; Abcam, UK; ab175473), goat anti-rabbit IgG H&L (Alexa Fluor 568; 1:1000; Abcam, UK; ab175471), goat anti-mouse IgG H&L (Alexa Fluor 647; 1:1000; Abcam, UK; ab150115), goat anti-rabbit IgG H&L (Alexa Fluor 647; 1:1000; Abcam, UK; ab150075), or goat anti-rat IgG H&L (Alexa Fluor 647; 1:1000; Abcam, UK; ab150159) for 9EG7 staining. Following three additional washes with DPBS, nuclei and F-actin were stained with NucBlue Fixed Cell Stain ReadyProbes reagent (Invitrogen, USA; 2831256) and ActinGreen 488 (Invitrogen, USA; 2647610), respectively, for 15 min at room temperature.

### Single-Cell Migration Tracking

Random cell migration was analyzed by time-lapse single-cell tracking. Cells were labeled with CellTracker Green CMFDA (Thermo Fisher Scientific, USA) at 5 μM for 30 min at room temperature and subsequently subjected to the indicated peptide treatments. For peptide-coated substrate experiments, migration imaging was initiated after 12 h of cell attachment. Time-lapse fluorescence images were acquired using a CellCyte live-cell imaging system at 6 min intervals for 2 h. Individual cell trajectories were reconstructed using TrackMate^45^ in Fiji, and migration velocity was quantified from the resulting trajectories.

For Ca²⁺ perturbation condition, peptide assemblies were prepared in the presence of 50 mM CaCl₂ and equilibrated under the same conditions used for the corresponding untreated assemblies. The Ca²⁺-treated peptide solutions were subsequently applied to the culture substrates and dried under vacuum overnight. Before cell seeding, the coated substrates were washed three times with DPBS for 20 min each to remove residual free Ca²⁺ and minimize direct effects of extracellular Ca²⁺ on the cells. Cell migration was then assessed using the same procedure described above.

For pharmacological inhibition experiments, cells were seeded on the indicated substrates and allowed to adhere for 11 h. For pharmacological treatment, cells were exposed to PF-573228 (10 μM; MedChemExpress, USA; HY-10461) or (−)-blebbistatin (20 μM; MedChemExpress, USA; HY-13441). An equivalent volume of DMSO was used as the vehicle control. After 1 h of treatment, time-lapse single-cell migration tracking was initiated and performed as described above.

### Post-Adhesion Soluble Peptide Treatment

HTR-8/SVneo cells were seeded on TC-treated culture dishes and allowed to establish adhesion for 12 h before peptide treatment. The culture medium was then replaced with DMEM containing 1% FBS and the indicated concentrations of peptide assemblies. Unless otherwise specified, cells were maintained in the presence of peptide for 12 h before subsequent single-cell migration tracking, confocal imaging, immunocytochemistry, or scanning electron microscopy. For assays requiring different treatment durations, including wound healing, Transwell migration, CCK-8, and Live/Dead assays, the corresponding incubation times are specified in the respective sections below.

### Scanning Electron Microscopy (SEM)

After 12 h peptide treatment, the culture medium was removed, and cells were washed three times with DPBS. Cells were fixed with 2.5% glutaraldehyde in 0.1 M cacodylate buffer for 30 min, followed by post-fixation with 1% OsO₄ (SPI Supplies, USA; 02604-AB) in 0.1 M cacodylate buffer for 30 min. Samples were then washed three times with Milli-Q water for 5 min each, dehydrated through a graded ethanol series, rinsed with tert-butanol (Innochem, China; KDHFT01), and freeze-dried for more than 12 h using a VTLG-10C lyophilizer (Yetuo, China). Before imaging, samples were sputter-coated with a 3 nm platinum layer using an EM ACE600 coating system (Leica, Germany). SEM images were acquired using a Gemini 300 field-emission scanning electron microscope (ZEISS, Germany).

### CCK-8 Cell Viability Assay

HTR-8/SVneo cells were seeded in 96-well plates (Merck, Germany; CLS3596) at a density of 3 × 10³ cells per well and allowed to adhere for 12 h at 37 °C in a humidified atmosphere containing 5% CO₂. The medium was then replaced with DMEM containing 1% FBS and the indicated concentrations of peptide, and cells were incubated for an additional 24, 48, or 72 h. Cell viability was assessed using a CCK-8 assay (Beyotime, China; C0046) according to the manufacturer’s instructions. Absorbance at 450 nm was measured using a Synergy H1 multimode microplate reader (BioTek, USA).

### Live/Dead Cell Viability Assay

HTR-8/SVneo cells were seeded in 96-well plates at a density of 3 × 10³ cells per well and allowed to adhere overnight. The medium was then replaced with DMEM containing 1% FBS and the indicated concentrations of peptide, and cells were treated for 72 h. After treatment, cells were stained with Calcein-AM and propidium iodide (PI) (Beyotime, China; C2015L) together with Hoechst 33342 (Beyotime, China; C1028) according to the manufacturer’s instructions and incubated at 37 °C for 30 min in the dark. Fluorescence images were acquired using a Nikon ECLIPSE Ts2 microscope (Nikon, Japan) with green, red, and blue fluorescence channels corresponding to live cells, dead cells, and nuclei, respectively. At least five randomly selected fields were acquired from each well, and live/dead cell fractions were quantified using ImageJ.

### Transwell Migration Assay

Transwell migration behavior was assessed using Transwell inserts with 8.0 μm pores. Serum-starved HTR-8/SVneo cells were seeded into the upper chamber at 4.5 × 10⁴ cells per insert in the presence or absence of the indicated peptide assemblies, while medium containing 20% FBS was placed in the lower chamber as a chemoattractant. After 24 h, non-migrated cells remaining on the upper surface of the membrane were removed. Cells that had migrated to the lower surface were fixed with 4% paraformaldehyde for 10 min, stained with Crystal Violet Staining Solution (Beyotime, China; C0121) for 10 min, thoroughly washed with water, imaged by microscopy, and quantified using ImageJ.

### Wound Healing Assay

HTR-8/SVneo cells were seeded in 96-well plates at a density of 5 × 10⁴ cells per well and cultured for 12 h to reach approximately 90% confluence. Uniform wounds were generated using an Essen Bioscience Wound Maker. After removal of detached cells by gentle washing, the medium was replaced with DMEM containing 1% FBS and the indicated concentrations of peptide assemblies. Wound closure was monitored by EVOS M7000. Images were acquired at defined intervals for 43 h, and migration was quantified as relative wound density using ImageJ.

### Western Blotting

Cells were lysed in RIPA buffer (Thermo Fisher Scientific, USA) supplemented with protease and phosphatase inhibitors (Thermo Fisher Scientific, USA) on ice for 30 min. Cell lysates were centrifuged at 15,000 × g for 30 min at 4 °C, and the supernatants were collected. Protein concentrations were determined using a BCA Protein Assay Kit (Thermo Fisher Scientific, USA). Equal amounts of protein (20–50 μg per lane) were separated by SDS-PAGE and transferred to PVDF membranes (Millipore, USA) using a Trans-Blot Turbo Transfer System (Bio-Rad, USA) at 25 V for 30 min. Membranes were blocked with blocking buffer (Beyotime, China; P0252) for 1 h at room temperature and incubated overnight at 4 °C with anti-phospho-myosin light chain 2 (Thr18/Ser19) antibody (1:1000; Cell Signaling Technology, USA; 3674S) or anti-GAPDH antibody (1:1000; Abcam, UK; ab8245). After three washes with TBST, membranes were incubated with the appropriate HRP-conjugated secondary antibodies, goat anti-rabbit IgG (1:5000; Invitrogen, USA; G21234) or goat anti-mouse IgG (1:5000; Invitrogen, USA; G21040), for 1 h at room temperature. Protein bands were visualized using enhanced chemiluminescence reagent (Thermo Fisher Scientific, USA; 34096) and imaged using a ChemiDoc imaging system (Bio-Rad, USA). Band intensities were quantified using ImageJ and normalized to GAPDH.

### Statistical Analysis

No statistical methods were used to predetermine sample size. Experiments were not randomized, and investigators were not blinded to group allocation during data collection or analysis. Unless otherwise specified, all experiments were independently repeated at least three times. The statistical tests used for each dataset are specified in the corresponding figure legends. Statistical analyses were performed using GraphPad Prism version 10.6.1 (GraphPad Software, USA). Data are presented as mean ± s.d., unless otherwise indicated. Adjustments for multiple comparisons were applied where appropriate, as specified in the figure legends. Exact p values are reported whenever applicable, and statistical significance was defined as p < 0.05.

## ASSOCIATED CONTENT

### Supporting Information

The following file is available free of charge. Additional experimental details for peptide characterization and molecular dynamics trajectory analysis; characterization of the scrambled peptide sFV (Figure S1); CD spectra of terminal-modified peptide variants (Figure S2); TEM characterization and width analysis of peptide assemblies (Figure S3); FTIR spectra of peptide assemblies (Figure S4); structural evolution, temporal robustness, and connectivity analyses of peptide assemblies in molecular dynamics simulations (Figures S5–S8); ligand exposure, mobility, and reproducibility analyses in molecular dynamics simulations (Figures S9–S10); characterization of peptide coatings by Congo red staining and AFM (Figure S11); cell attachment to peptide-coated substrates and β1-integrin-blocking experiments (Figures S12–S13); cell morphology, spreading, and integrin β1 activation after initial attachment (Figures S14–S15); paxillin and activated integrin β1 immunofluorescence (Figure S16); effects of Ca²⁺ perturbation on peptide-regulated cell migration (Figure S17); cell morphology, spreading, paxillin organization, and phosphorylated myosin light chain after prolonged attachment to peptide-coated substrates (Figures S18–S20); effects of blebbistatin and FAK inhibition on cell migration (Figure S21); cell viability and concentration-dependent migration under post-adhesion peptide-treatment conditions (Figures S22–S23); SEM characterization of cell–assembly interactions (Figure S24); cell spreading after post-adhesion treatment with FV and FV-NH₂ (Figure S25); and HPLC, LC–MS, and ^1H NMR characterization of the synthesized peptides (Attachments S1–S8) (PDF).

## AUTHOR INFORMATION

### Author Contributions

J.G., Z.Y., Y.Z., and X.H. conceived the study and designed the experiments. Z.S., Y.L., and X.H. performed FTIR spectroscopy, thermal perturbation experiments, preparation and characterization of peptide-coated substrates, Congo Red staining, AFM imaging, cell attachment assays, single-cell migration assays, wound-healing assays, Transwell migration assays, immunocytochemistry and confocal microscopy, Western blotting, and SEM imaging. Z.Z. and X.L. performed CD spectroscopy, TEM characterization, and Nile Red fluorescence assays. Q.Z. conducted NMR spectroscopy. W.H. and X.H. performed molecular dynamics simulations and trajectory analyses. Y.C., J.G., Z.Y., and Y.Z. advised the study. X.H. wrote the manuscript.

## Notes

The authors declare the following competing financial interest(s): X.H. and Y.Z. are inventors on a patent application related to the data presented in this manuscript.

## ACKNOWLEDGMENT

The authors thank Dr. Ting Liang for assistance with HPLC and mass spectrometry measurements, and Junchao Zhi and Zhen Gao for assistance with cell culture, all at the Dongguan Institute of Materials Science and Technology, Chinese Academy of Sciences. The authors also thank Dr. Xiaofang Zhang for providing training in TEM imaging; Huanglei Yu, Xiu Li, and Yupeng He for providing training in AFM sample preparation and measurements; Jing Sun for assistance with FTIR measurements; and Jinsheng Liu for assistance with SEM measurements, all at Songshan Lake Materials Laboratory.

DMSO: dimethyl sulfoxide
FAK: Focal Adhesion Kinase
LC-MS: liquid chromatography-mass spectrometry
LINCS: LINear Constraint Solver
NVT: canonical ensemble
NPT: isothermal-isobaric ensemble
CCK-8: Cell Counting Kit-8

